# Acute toxicity of the plant volatile indole depends on herbivore specialization

**DOI:** 10.1101/784165

**Authors:** Abhinav K. Maurya, Rakhi C. Patel, Christopher J. Frost

**Affiliations:** Department of Biology, University of Louisville, Louisville, Kentucky, 40292, USA

**Keywords:** Caterpillars, Green leaf volatiles (GLVs), Herbivore-induced plant volatiles (HIPVs), Host Range, Indole, Pest, Toxicity, Specialist vs Generalist herbivore

## Abstract

Herbivore-induced plant volatiles (HIPVs) provide direct benefits to plants as antimicrobials and herbivore repellents, but their potential as direct toxins to herbivores is unclear. Here we assayed the larvicidal activity of six common HIPVs from three different biochemical pathways and tested the hypothesis that the larvicidal activity of HIPVs is related to the host specialization of the insect pest. We first assessed β-caryophyllene, linalool, *z*-3-hexenyl acetate, *z*-3-hexenol, *e*-2-hexenol, and indole against the beet armyworm (*Spodoptera exigua*), and found that indole was 7-fold more toxic compared to the other volatiles when incorporated into diet. Then, we tested the larvicidal activity of indole against six common, destructive pest caterpillars with varying host ranges. Consistent with our hypothesis, indole toxicity varied with caterpillar host range: indole toxicity was seven-fold higher in more specialized insect species relative to generalist insect species. That said, the LC_50_ of indole was comparable to other reported anti-herbivore agents even against the generalist caterpillars. Yet, indole in headspace had neither larvicidal nor ovicidal activity on any caterpillar species tested. These results support a key ecological precept regarding tradeoffs between host specialization and chemical detoxification, and also indicate that indole functions as a direct defense against herbivores that could be potentially useful in integrated pest management strategies.

**Key message:**

- We measured the direct toxicity of six common HIPVs against the beet armyworm.
- Indole was the most toxic HIPV against the beet armyworm.
- We determined the toxicity of indole against six different pest caterpillar species.
- Toxicity of indole was associated with the host preference of the insect species.
- Indole exposure in headspace had no effect on egg hatching or caterpillar survival.
- Indole has the potential to be developed as an insecticide against crop pests.

**Author Contribution Statement:** CJF and AKM designed research. AKM and RCP conducted experiments. AKM and CJF analyzed data and wrote the manuscript. All authors read and approved the manuscript.

## Introduction

Plants produce a remarkable variety of volatile organic compounds (VOCs) that can affect the behavior of pollinators (Schiestl and Ayasse 2001; Schiestl et al. 1999), seed dispersers (Valenta et al. 2017), and herbivores (Agrawal 2001; Vickers et al. 2009). A subclass of VOCs is herbivore-induced plant volatiles (HIPVs), which plants release in response to herbivore attack. HIPVs are typically a blend of compounds derived from multiple biosynthetic pathways including terpenes, fatty acid derivitatives, and shikimate derivitatives. HIPVs confer both indirect and direct defense benefits (Hare 2011), act as priming cues that activate plant defenses and reduce herbivory (Erb et al. 2015; Frost et al. 2007; Frost et al. 2008a; Frost et al. 2008b; Heil and Bueno 2007), attract natural enemies (Birkett et al. 2003; Dicke 1986; Dicke and Sabelis 1988; Schnee et al. 2006; Turlings et al. 1995; Turlings et al. 1990; Zhu and Park 2005), and can even render herbivores susceptible to entomopathogens (Gasmi et al. 2018). HIPVs also have direct defense benefits to the plants that produce them, including protecting plants from microbial and pathogen infections (Shiojiri et al. 2006; Yi et al. 2009), deterring herbivory (Beale et al. 2006; Bernasconi et al. 1998; Heil 2004; Sandra et al. 2014) and oviposition (Kessler and Baldwin 2001; Veyrat et al. 2016; Zakir et al. 2013), and reducing caterpillar growth (von Mérey et al. 2013) and food consumption (Veyrat et al. 2016) after just HIPV exposure.

The hypothesis that HIPVs directly affect insect herbivores fecundity is not new (Pichersky and Gershenzon 2002; Unsicker et al. 2009), but the direct larvicidal or ovicidal efficacy of HIPVs on insect herbivores is poorly understood. This is due in part to the fact that the consideration of diverse phytochemicals acting as a selective pressures driving insect pest feeding strategies has largely excluded volatile consitutents (Endara et al. 2017; Feeny 1976; Howard V. C. and Bradford A. H. 2003). The vast majority of insects have evolved host range specialization, feeding on only one or a few closely related species (Forister et al. 2015), while a minority of insect herbivore species has a more generalist host range. Evolutionary theory predicts that phytochemicals that are widespread among different plant taxa will be less toxic to generalist insects compared to specialists (Howard V. C. and Bradford A. H. 2003). HIPVs tend to be common across plant taxa, and some HIPVs can be pre-synthesized, stored in specialized cells in their original or conjugate forms in various types of plant tissues (Baldwin 2010; Monson et al. 2012; Ormeño et al. 2011; Sugimoto et al. 2015; Tominaga and Dubourdieu 2000), and released when herbivory disrupts cellular storage compartments (Niinemets et al. 2013). Insect pests must therefore cope with potential toxic effects of HIPVs by either direct ingestion or headspace exposure.

The insect order Lepidoptera (butterflies and moths) contains many of the major agricultural pests that cause significant damage and economic loss of food crops worldwide (Vreysen et al. 2016). Known lepidopteran pests include both feeding specialists and generalists. To combat these pests, potent and toxic synthetic chemicals are frequently used in current agricultural systems (Cordero et al. 2006; Ecobichon 2001; Pimentel 1996). However, these insecticides can have detrimental health effects on humans and non-target organisms (Cimino et al. 2016; Hahn et al. 2015; Mulé et al. 2017; Tingle et al. 2003), and the evolution of resistance by insect pests against commonly used insecticides is well documented (Brown 1958; Sparks and Nauen 2015). Plants produce a variety of chemicals with insecticidal activity, and even some VOCs are directly toxic against invertebrates (Hubert et al. 2008; Laquale et al. 2018; Lee et al. 1999; Zhao et al. 2017). Moreover, blends of plant essential oils containing major constituents of HIPV blends (Maffei et al. 2011) are known ovicidals and larvicidals against lepidopteran pests (Bakkali et al. 2008; El-Zaeddi et al. 2016; Isman 2016; Mossa 2016). Although the potential toxicity of individual HIPVs against lepidopteran pests is limited, investigating the larvicidal and ovicidal activity of common individual HIPVs may add to our arsenal of chemical-mediated pest control options in agriculture systems.

Here, we evaluated the hypothesis that HIPVs are acutely toxic to insect herbivores. Plant volatiles may affect herbivores either as a constituent of ingested leaf tissues or through air contact alone (Veyrat et al. 2016), so we conducted dose-response assays with HIPVs either in headspace alone or infused directly into diet. We specifically selected six HIPVs that represented the three major biochemical pathways: terpenes, green leaf volatiles (GLVs) derived from the lipoxygenase pathway, and indole (Heil 2014). Indole is an aromatic, bicyclic, amino acid precursor from the shikimate pathway. Volatile indole is emitted from a variety of eukaryotes and prokaryotes (Lee et al. 2015), and recently indole has been implicated in plant defense priming and direct defense against insects (Erb et al. 2015).

First, we tested the larvicidal activity the six individual HIPVs against a common lepidopteran herbivore pest beet-armyworm (*Spodoptera exigua*). We used the beet armyworm (*S. exigua*) in our first experiments because it is destructive generalist agricultural pest (Liburd et al. 2000; Pearson 1983) that has developed resistance against chemical insecticides (Brewer et al. 1990; Che et al. 2013), and is also a model herbivore in HIPV-mediated direct and indirect plant defense studies (Christensen et al. 2013; Engelberth et al. 2004; Huffaker et al. 2013; Jurriaan et al. 2007; Schmelz et al. 2003). As non-volatile terpenes and phenylpropanoids are known direct defenses (Moghaddam and Mehdizadeh 2017), we predicted that indole and the terpenes would be relatively more toxic than the GLVs. Results from this experiment led us to focus our work specifically on indole. We tested the larvicidal activity of indole on six agriculturally important caterpillar species with different host ranges. Because indole is produced by a wide range of plant species (Cna’ani et al. 2018; Lee et al. 2015), we hypothesized that indole toxicity would increase with herbivore host specialization. That is, specialists would be more sensitive to indole than would be generalists. Lastly, we tested the ovicidal effect of indole. Since HIPVs provide indirect defenses by attracting egg predators (Fatouros et al. 2008), we predicted that indole would provide a direct defense benefit by reducing egg hatching success.

## Materials and Methods

### Plant volatiles

We used six common, commercially available HIPVs belonging to different biosynthetic pathways. Three compounds were GLVs derived from the lypoxygenase pathway: *cis*-3-hexenol (97%) (CAS: 928-96-1; TCI America), *cis*-3-hexenyl acetate (99%) (CAS: 3681-71-8; TCI America), and *trans*-2-hexenal 98%) (Sigma-Aldrich). Two terpene representatives were the sesquiterpene β-caryophyllene 97% (CAS: 87-44-5; MP Biomedicals) and the monoterpene linalool (97%) (CAS: 78-70-6; Alfa Aesar). Finally, we tested the nitrogen-containing compound indole (97%) (CAS: 120-72-9; TCI America) that derives from the shikimic acid pathway.

### Experimental insects

For our experiments, we selected six common pest herbivores that are known to cause severe economic losses. We chose four generalists: beet armyworm (*Spodoptera exigua*) (Capinera 1999a; Greenberg et al. 2001), fall armyworm (*Spodoptera frugiperda*) (CABI; Capinera 1999c), cotton bollworm (*Helicoverpa zea*) (CABI; Martin et al. 1976), and tobacco budworm (*Heliothis virescens*) (Capinera 2001; Harding 1976; Martin et al. 1976). We selected the cabbage looper (*Trichoplusia ni*), a generalist that has a strong host preference for the mustard family (Brassicaceae) (Capinera 1999b; Hoo et al. 1984; Martin et al. 1976). Finally, we selected velvetbean caterpillar (*Anticarsia gemmatalis*), a specialist on legumes (Slansky 1993) (Waters and Barfield 1989). Eggs or egg masses of these species were obtained from Benzon Research Inc. USA (Permit #P526P-16-02563 to CJF). Eggs were immediately transferred to 2-ounce diet cups for hatching. The diet cups were maintained on shelving in a climate-controlled room at 24°C-27°C until egg hatched, and 1^st^ instar larvae were used within 24 h of hatching for all experiments.

### Preparation of test diets for feeding bioassays

Larvicidal effects of HIPVs against *S. exigua* were tested at five different concentrations 1, 2.5, 3.75, 5, and 10 mg/ml or μl/ml in feeding and headspace bioassays. Previous work establishing the LC_50_ of *trans*-2-hexenal against five species of stored-product beetles (Hubert et al. 2008) was used as a starting point for initial test concentrations in our study, and preliminary assays suggested this range would be adequate to determine LC_50_ for the HIPVs. However, the relatively high larvicidal activity of indole in our initial experiments caused us to also include lower concentrations of indole ranging from 0.005 to 1 mg/ml. All test diets were prepared 12 h prior to starting an experiment. Artificial diet powder (Southland Products Incorporated, Arkansas, USA) was prepared per manufactures instructions and aliquoted into 50-ml centrifuge tubes. Prior to the diet solidifying, an appropriate amount of an individual HIPV was added, and the tube was vortexed thoroughly to mix each test compound in the diet. Control diets were prepared similarly but without any HIPV added. After solidifying at room temperature, the diet was cut into disc-shaped pieces (10 mm diameter, 5 mm height, ca. 400 mg) using a 10-cm long cork borer. Each experimental cup received one piece of artificial diet.

### Preparation of volatile dispenser for headspace bioassays

Experimental amounts of *cis*-3-hexenol, c*is*-3-hexenyl acetate, β-caryophyllene, linalool, *trans*-2-hexenal, and indole was added into a 2.0ml amber glass vial (Agilent Technologies) with 1 mg of glass wool (Fig. 1). Control dispensers had only glass wool without any volatile (Erb et al. 2015). The amber vials with HIPVs were sealed with a rubber septum and connected to the diet cup by piercing the diet cup and amber vial rubber septum with an 18-gauge needle (inner diameter 0.83 mm). This design was similar to previous work in which dispensers containing 20 mg of indole were pierced with a 1 μL micro-pipette (inner diameter 0.2mm) (Ye et al. 2019).

**Fig. 1.**
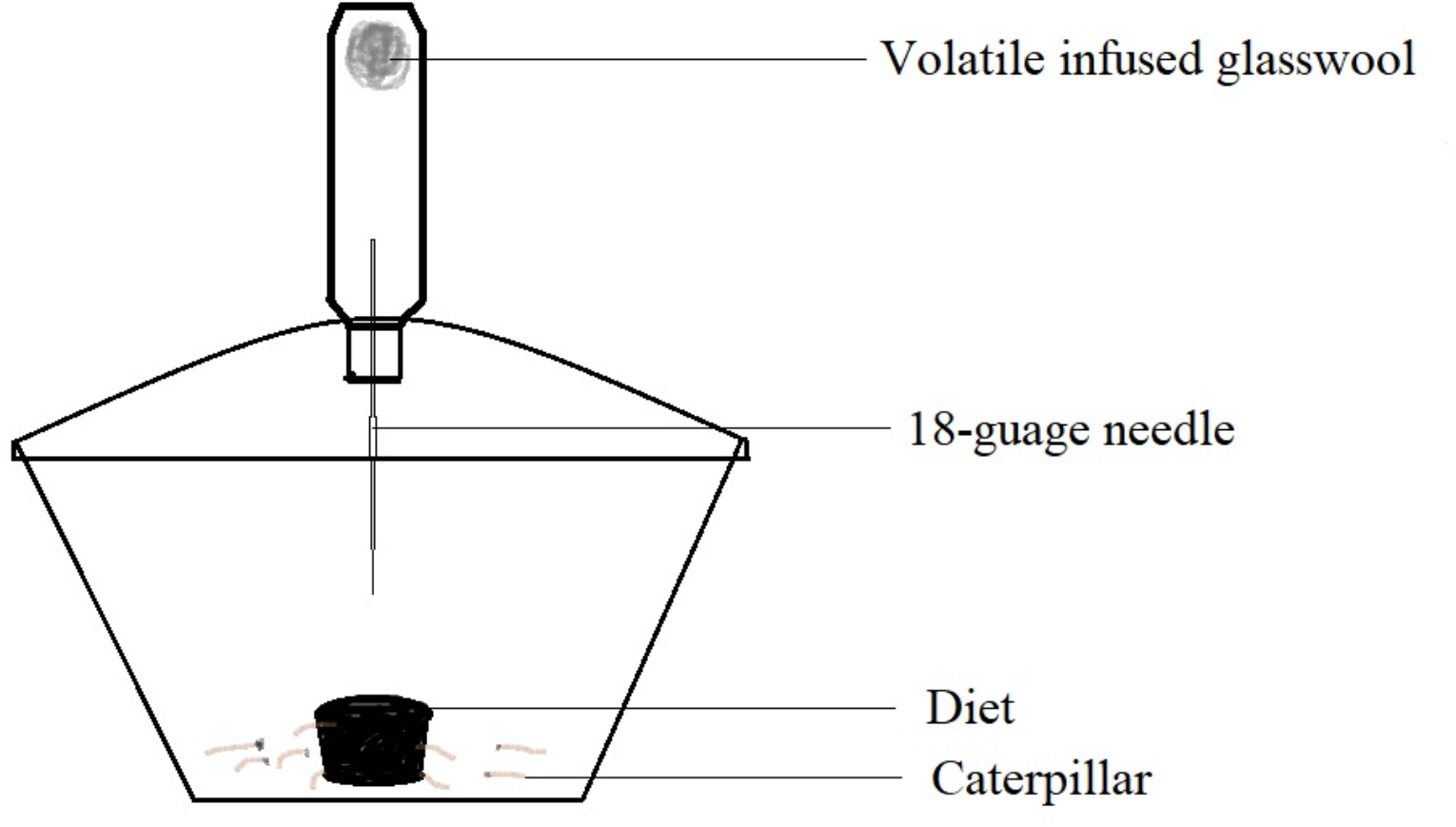
Volatile delivery system for headspace bioassay.

### Test of toxicity of plant volatiles against caterpillars

First instar larvae were used for testing the toxicity of plant volatiles because the first instar is the most sensitive stage to secondary plant chemicals (Zalucki et al. 2002). For feeding bioassays, ten first instar larvae were placed in a 2 oz diet cup containing either volatile infused diet or control diet. The cup was the experimental unit of replication. Headspace bioassays were conducted similarly, except that all diets were control (no HIPVs) and the diet cups were connected to a dispenser that contained a specific HIPV or no HIPV (control). Each experimental group had 5-10 replicate cups. The percent survival at 24 h was determined for each replicate.

### Effects of headspace indole on *S. exigua* and *T. ni* egg hatching rates

For egg hatching assays, we specifically selected caterpillar species most susceptible and tolerant to indole in feeding bioassays. The inhibitory effect of indole on the beet armyworm (*S. exigua*) and the cabbage looper (*T. ni*) egg hatching was measured in a headspace bioassay. *S. exigua* and *T. ni* eggs were transferred to diet cups that were connected to volatile dispensers containing different concentrations of indole: 0, 0.1, 0.25, 0.5, 1, 2.5, 5, 10, 15 and 20 mg. Each concentration of indole had 5 replicate diet cups. The percent hatch of the eggs was measured at 96 h after exposure and compared to controls without indole.

### Statistical analyses

All analyses were conducted in R version 3.4.1 (R 2018) using RStudio version 1.0.153 (Wickham 2011). For calculating the median lethal concentration (LC_50_), the mortality rates of caterpillar larvae after 24 hours of VOC exposure were regressed against indole concentration using quadratic logistic regression (glm function), and the median lethal concentration (LC_50_) was calculated using the fitted function in quadratic logistic regression. We used this model because it produced the best fit based on AIC values. Figures were plotted with GraphPad Prism version 8.0.0 (San Diego, California USA) and ggplot2 in R Studio (Wickham 2011).

## Results

### HIPV toxicity against *S. exigua*

Indole caused highest larval mortality of all six HIPVs tested (Fig. 2A). With the exception of β-caryophyllene, all HIPVs that were directly consumed in dietcaused complete mortality at some concentration tested (Fig. 2A). In contrast, no HIPV showed any toxicity to *S.exigua* when administered in headspace alone at any concentration (Fig. 2B). Based on LC_50_ values, indole was more than 7 times more larvicidal than was the second most potent volatile (linalool) among all the HIPVs tested against *S. exigua* (indole LC_50_ = 0.35 mg/ml; linalool LC_50_ =2.59 μl/ml) (Fig. 3). GLVs were relatively less toxic: *cis*-3-hexenol (LC_50_ = 3.32 μl/ml diet), *cis*-3-hexenyl acetate (LC_50_ =4.61 μl/ml diet) and *cis*-3-hexenal (LC_50_ =4.85 μl/ml diet) (Fig. 3). β-caryophyllene was neither toxic in diet nor headspace against *S.exigua* caterpillars at any of the tested concentrations. Mortality of *S. exigua* was negligible in the control group.

**Fig. 2.**
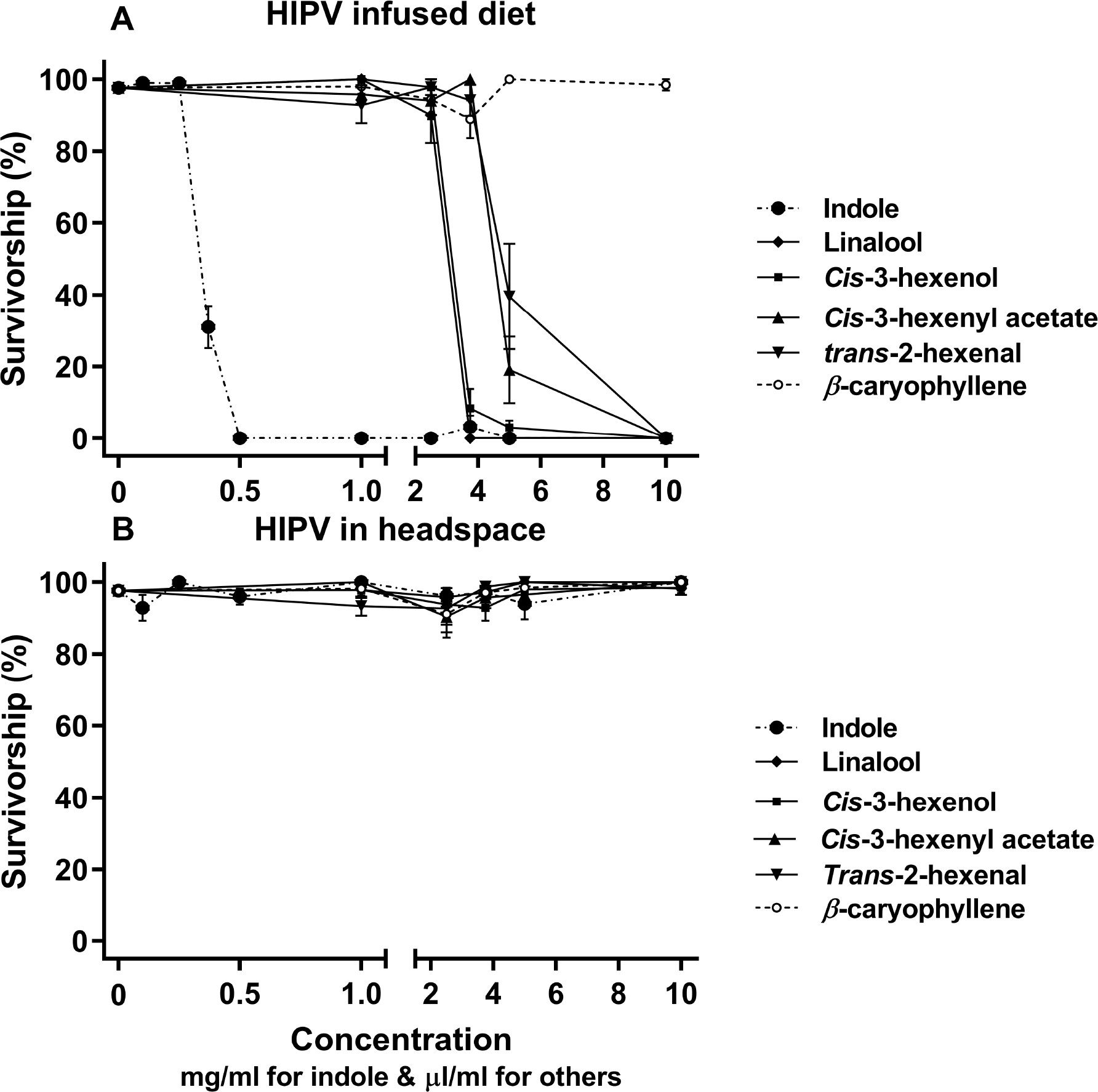
Direct toxicity of plant volatiles at concentrations on the survival of *S. exigua* in feeding bioassays (A) and headspace bioassay (B). Values at each concentration represent the mean of five-ten biological replicates ± 1SEM.

**Fig. 3.**
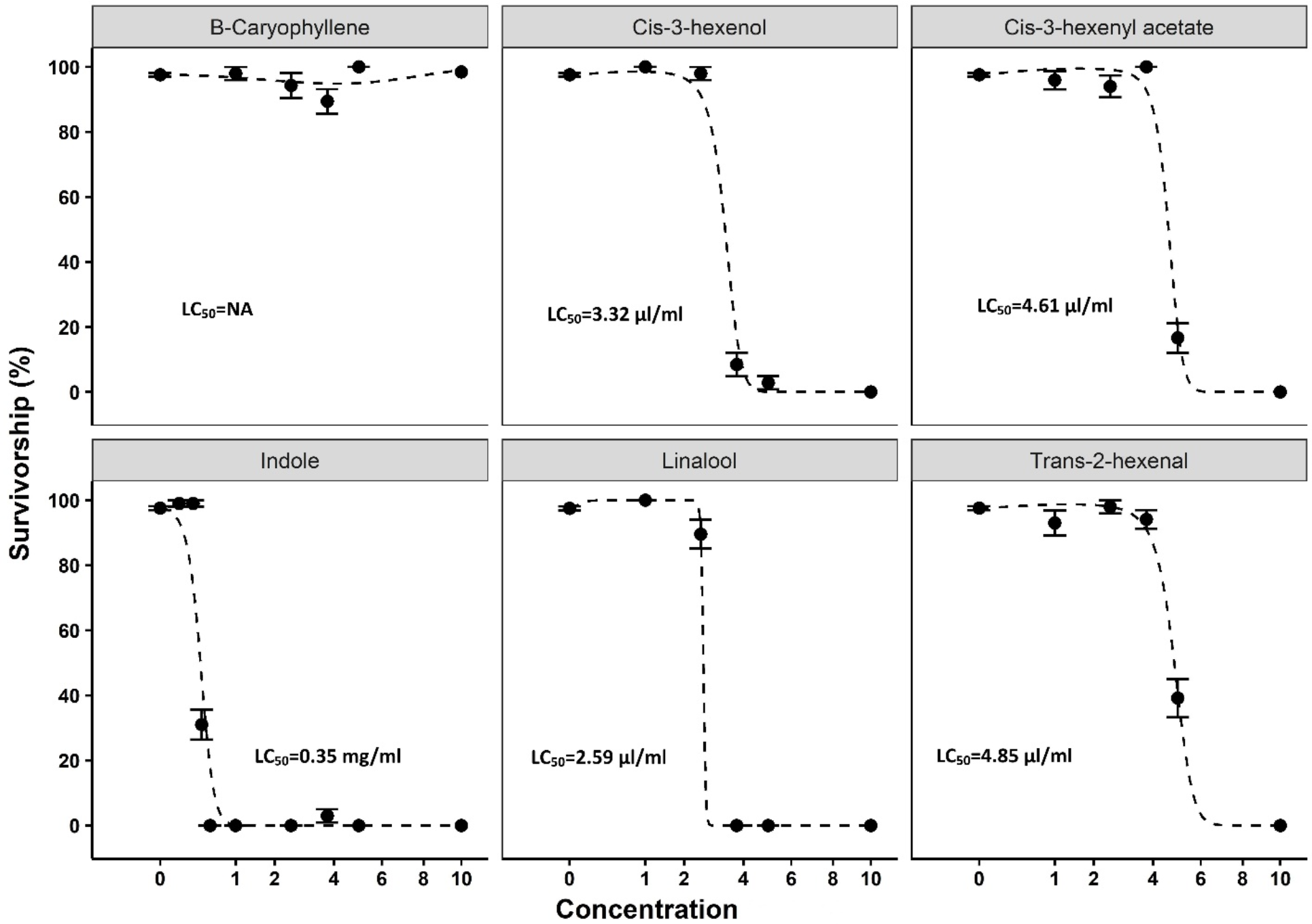
LC_50_ of individual plant volatiles on *S. exigua* caterpillars in feeding bioassay. Graphs for each volatile refer to fitted values based on quadratic logistic regression. LC50 represent lethal concentration causing 50 percent mortality. Data are reproduced individually from Fig.2A for easy visualization.

### Indole toxicity relative to caterpillar host range

The caterpillar species with restricted host ranges were more sensitive to indole than were the generalist caterpillars. The LC_50_ of indole for the four widely generalist pests ranged from 0.18-0.35 mg/ml diet (Fig 4). Specifically, the LC_50_ of indole was 0.35 mg/ml for *S.exigua* (Fig. 4A), 0.29 mg/ml for *S. frugiperda* (Fig. 4B), 0.27 mg/ml for *H. zea*. (Fig. 4C), and 0.18 mg/ml for *H. virescens* (Fig. 4D). In contrast, the LC_50_ of indole was 0.05 mg/ml for both the velvetbean caterpillar (*A. gemmatalis*) and the cabbage looper (*T.ni*) (Fig. 4E,F). That is, indole was 3.6-7.0 times more toxic for the two more specialized caterpillars than it was for the four generalists.

**Fig. 4.**
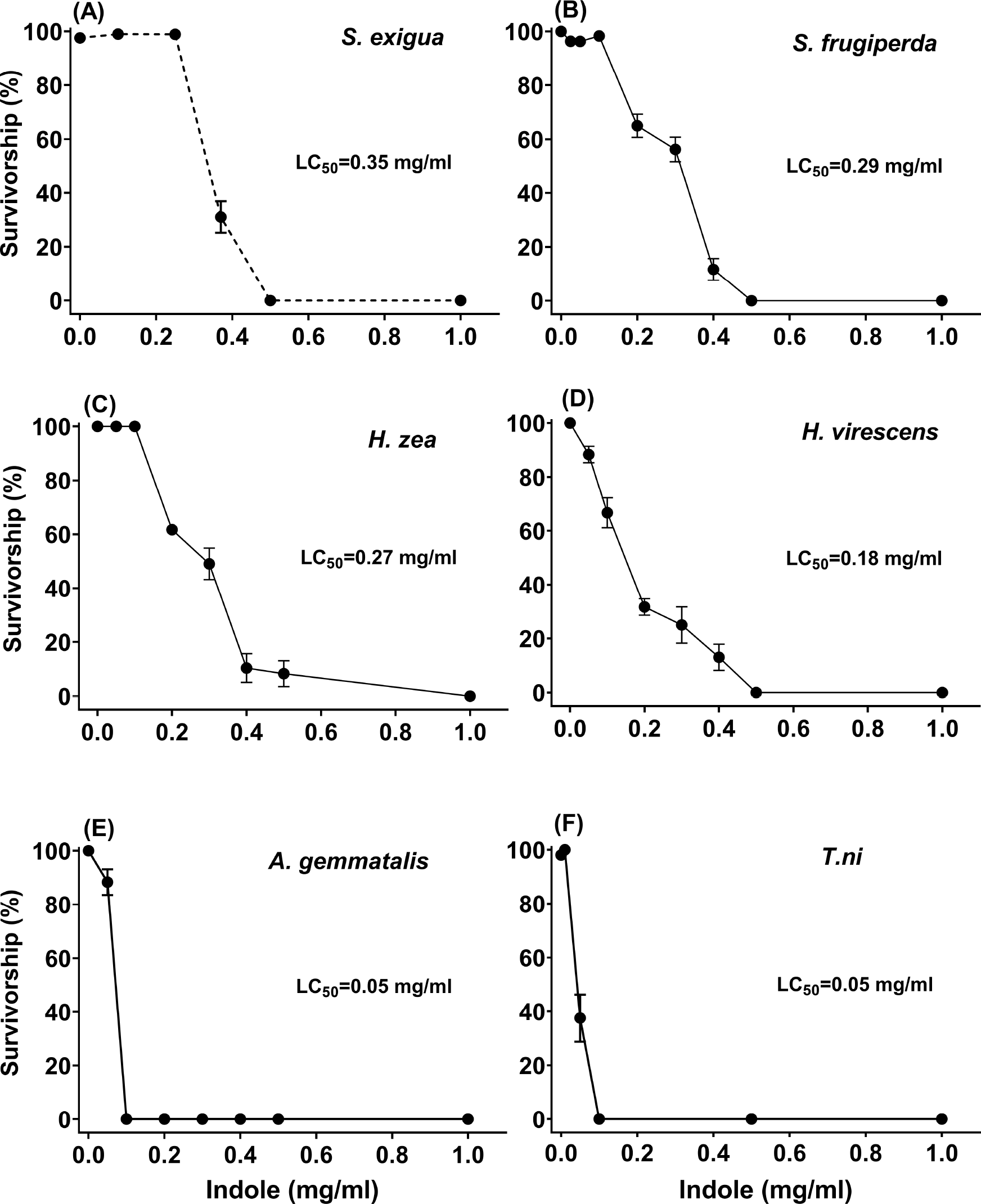
Direct toxicity of indole on the survival of five different caterpillar species in feeding bioassays. Values at each concentration represent the mean of five-ten biological replicates ± 1SEM. *S. exigua* data are reproduced (dashed lines) from Fig. 2 to aid in comparison.

### Larvicidal and ovicidal activity of volatile indole in headspace

Consistent with our first experiment using *S.exigua* (Fig. 2B), indole present only in headspace had no effect on *T. ni* caterpillar mortality (Fig.5). Furthermore, there was no inhibitory effect of any tested concentration of indole on egg hatching success of either *S. exigua* or *T. ni* caterpillars (Fig. 6A,B). Because these two caterpillar species were the most (*S.exigua*) and least (*T.ni*) sensitive to indole, we did not conduct headspace bioassays for mortality or egg hatching on the other caterpillars.

**Fig. 5.**
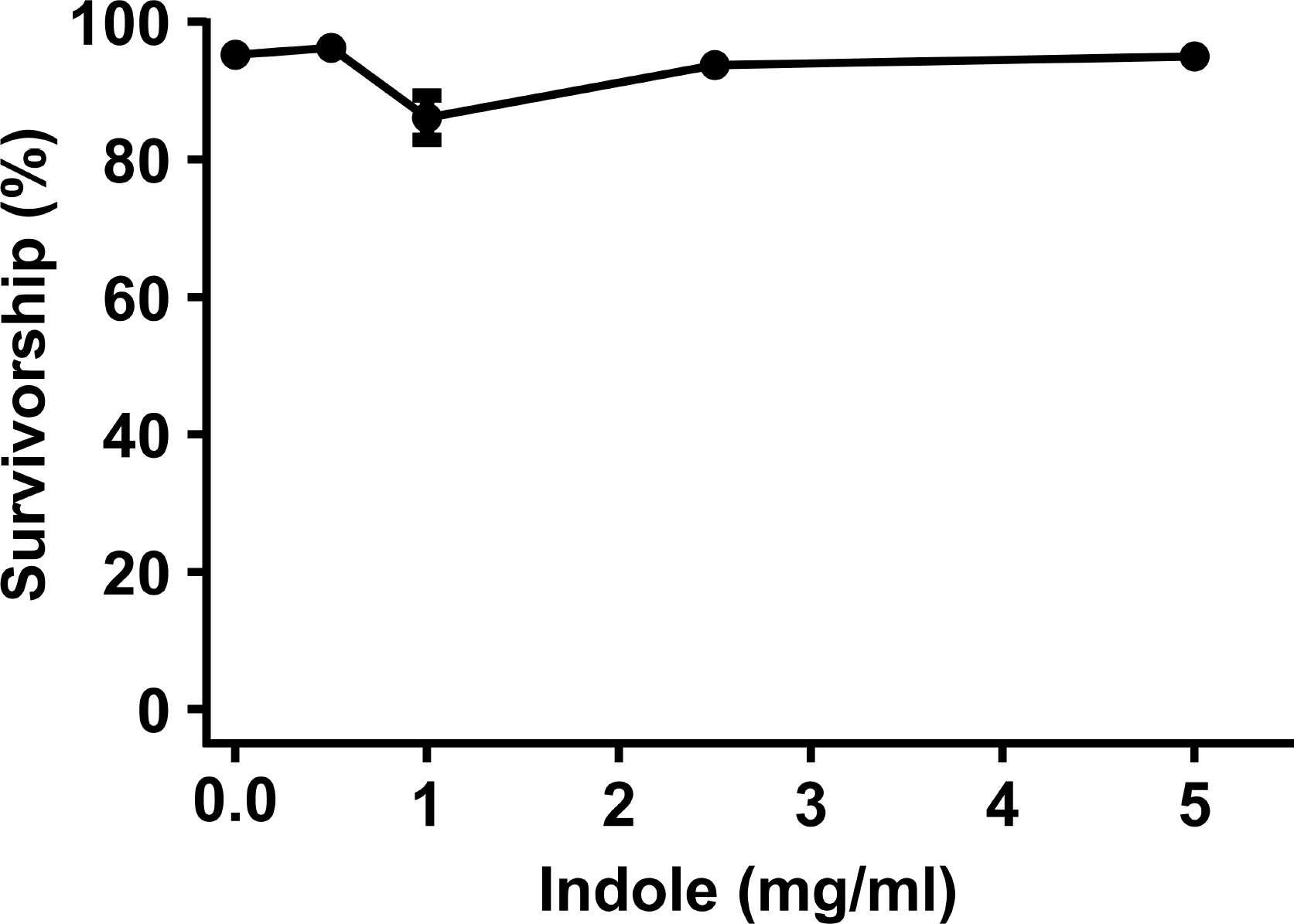
Effect of varying concentrations of indole on the survival of *T. ni* in headspace bioassays. Values at each concentration represent the mean of five-ten biological replicates ± 1SEM.

**Fig. 6.**
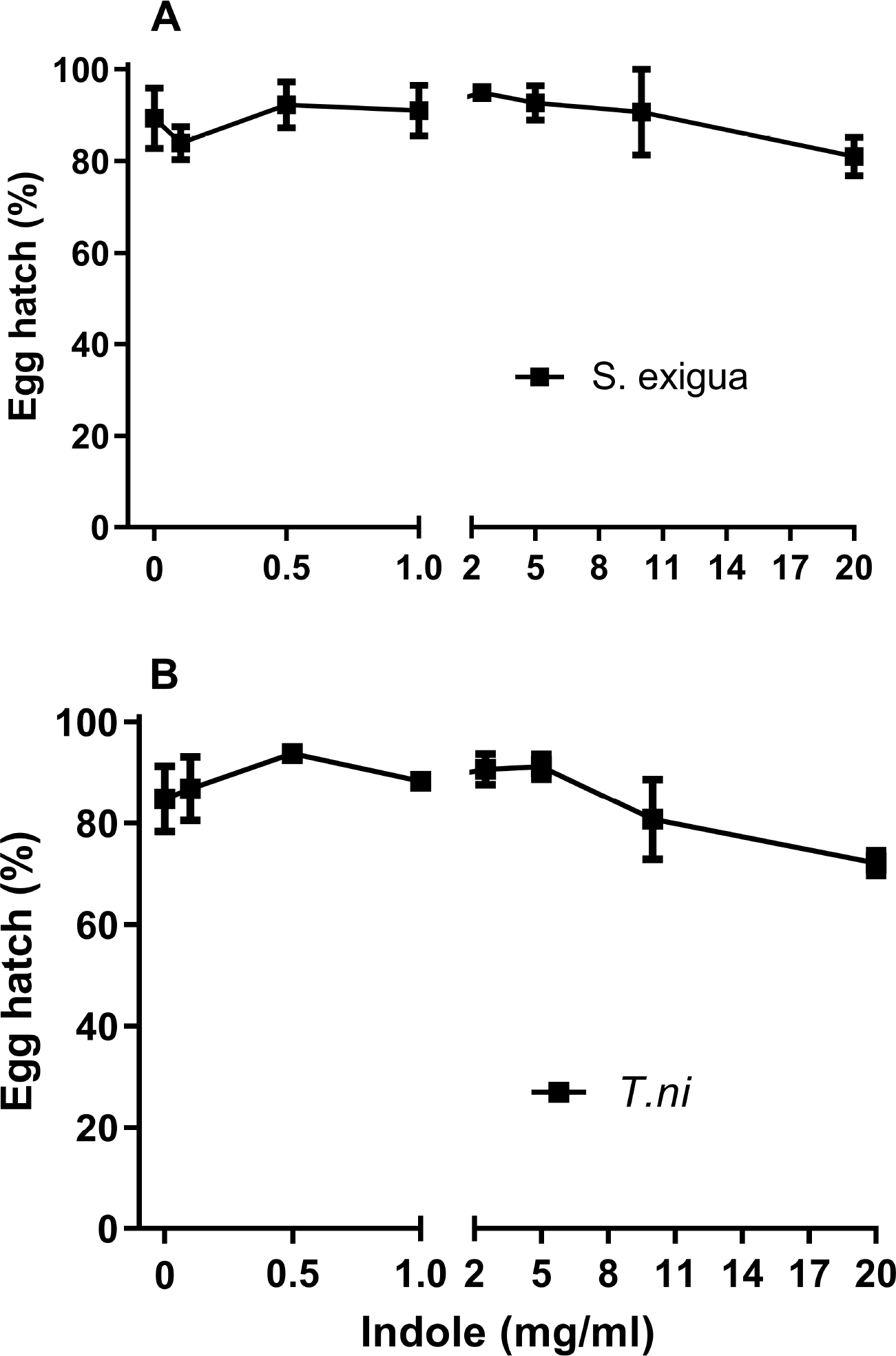
Effect of varying concentrations of indole on percent egg hatch of *S.exigua* (A) and *T. ni* (B). Values represent mean of five biological replicate ± 1SEM.

## Discussion

We have demonstrated that (1) HIPVs have direct larvicidal activity in an HIPV-specific manner, (2) the larvicidal effect of HIPVs is largely constrained to direct consumption by caterpillars as opposed to volatile exposure alone, and (3) the larvicidal effect of the common volatile indole depends on the degree of host specialization of the caterpillar species. To our knowledge, this is the first study to report LC_50_ values for HIPVs against lepidopteran pests, which was a key goal of this study. In supporting our prediction that larvicidal activity would vary among HIPVs, we found that indole and, to a lesser extent, GLVs and volatile terpenes, had direct larvicidal activity against the beet armyworm, a destructive pest with a wide host range. The larvicidal effect of some GLVs and terpenes has been reported against stored-pest beetles (Hubert et al. 2008) and aphids (Sadeghi et al. 2009), and our results are comparable. For example, the LC_50_ values we obtained for the three GLVs tested, *cis*-3-hexenol (3.32 μl/ml), *cis*-3-hexenyl acetate (4.61 μl/ml), and *trans*-2-hexenal (4.85 μl/ml) (Figure 3), are similar to those reported against stored pest beetles (0.6-3.32 mg/g) (Hubert et al. 2008). Similarly, the larvicidal activity of the terpene linalool against *S.exigua* in our study (LC_50_ of 2.59 μl/ml) is comparable to previous work testing linalool against the European corn borer (Lee et al. 1999). This suggests that the LC_50_ values we obtained may be broadly applicable across insect taxa.

A key finding is that indole is considerably more toxic than the other HIPVs tested. In fact, the LC_50_ of indole (50-350 μg/ml) is comparable to other natural toxicants such various strains of *Bacillus thuringiensis* (LC_50_ = 63.0-153.0 μg/ml) (Moar et al. 1989) and purified Cry1 protein from *B.thuringiensis* (LC_50_=1-870 μg/g) (Ali et al. 2006; Niu et al. 2013). Moreover, the LC_50_ of commercial *B. thuringiensis* DiPel ES (LC_50_= 2 μg/g)(Liao et al. 2002) and the synthetic insecticide lambda-cyhalothrin (LC_50_=5.27 μg/ml)(Hardke et al. 2011) are close to LC_50_ obtained for indole at 24h in our experiments. While the LC_50_ values of tested HIPVs in this study are higher than emission rates observed in nature (Allmann et al. 2013; Degen et al. 2012), they are likely representative of what might be stored within leaf tissues (Loreto et al. 1998; Loreto et al. 2000; Niinemets et al. 2004) and what insect herbivores may realistically encounter in their natural diets. In other words, our results have both ecological and practical relevance.

In our study, indole in headspace alone did not affect caterpillar survival or egg hatching. In fact, none of the volatiles tested showed any effect when present in headspace alone (Fig. 2B). In previous work, volatile indole reduced survival of the generalist herbivore *S. littoralis* by ~10% in neonates and ~6% in 1^st^ instar (Veyrat et al. 2016). This study also noted decreases in food consumption and, surprisingly, increases in larval weigh under indole treatments (Veyrat et al. 2016). While we did not find similar effects with *S.exigua*, our study was focused on larvicidal activity as reflected in LC_50_, which is fundamentally different metric. So, it appears that volatile exposure may exert non-lethal influences on caterpillar fecundity. That said, exposure of eggs to indole in headspace also had no effect on hatching success of either *S.exigua* or *T.ni*. (Figure 6), which is consistent with previous work (Veyrat et al. 2016). Plants volatiles can clearly have repellent effects on insect pests (Beale et al. 2006; Bernasconi et al. 1998; Heil 2004; Sandra et al. 2014), but our study indicates that larvicidal efficacy of HIPVs depends on their direct consumption by herbivores.

The larvicidal activity of indole varied with caterpillar host range in a manner that supported our initial prediction. A long-standing hypothesis is that generalist herbivores are well-equipped to detoxify wide array of common phytochemicals (Agrawal and Ali 2012; Krieger et al. 1971), whereas specialist herbivores are more tolerant to compounds specific to their host range but sensitive to more common phytochemicals (Whittaker and Feeny 1971). In other words, there is a tradeoff between chemical detoxification and host specialization. Since free indole is produced in some eukaryotes and prokaryotes (Lee et al. 2015) and is a common HIPV (Cna’ani et al. 2018; Frost et al. 2007), we predicted that it would be relatively toxic to specialist caterpillars compared to generalist caterpillars. Indeed, larvicidal activity of indole was approximately seven times lower for specialist *A. gemmatalis* compared to generalist *S. exigua* and, in general, all the generalist species were tolerant to indole relative to the specialist. The exception was *T. ni*, a pest that displays clear feeding preferences for Brassicaceae species (Rivera-Vega et al. 2017), which had an LC_50_ to indole approximately same as the specialist *A. gemmatalis* (LC_50_=0.05 mg/ml of diet) (Figure 4). While we only used six caterpillar species in our study, our results support ecological theory related to herbivore specialization.

Indole, and other HIPVs, may be useful insecticides in agricultural systems. More than 500 insect pest species have developed documented resistance to chemical insecticides (Georghiou 1990). Agriculturally destructive caterpillars are particularly capable of developing such resistance (Ahmad et al. 2008; Che et al. 2013; Elzen 1997; Hardee et al. 2001; McEwen and Splittstoesser 1970; Yu et al. 2003). Plant secondary metabolites are among the alternatives to synthetic insecticides in pest management, while also potentially avoiding or ameliorating negative impacts on beneficial organisms. Our study identifies that indole, and potentially other plant-derived volatiles, may be important additions to the arsenal of chemical defenses in pest management. Due to its toxicity, indole might be particularly useful as part of integrated pest management strategy for both generalist and specialist caterpillars given its larvicidal effect is similar or stronger than to some commercial pesticides and potent biopesticides like the Cry1F *bacillus thuringiensis* protein. Even the larvicidal activity of the GLVs and linalool against *S.exigua* in our study is approximately the same as reported for other pests. Therefore, our results indicate that HIPVs warrant attention as chemical components of control strategies against insect pests.

## Acknowledgments

This work was supported by a National Science Foundation grant IOS-1656625 to CJF. We thank Dr. Gary Cobbs for help with statistical analysis, members of the Frost Lab and Steve Yanoviak for comments on previous versions of the manuscript, and Rachel Haslem for technical support.

## References

Agrawal AA (2001) Phenotypic plasticity in the interactions and evolution of species. Science 294:321–326

Agrawal Aa, Ali JG (2012) Specialist versus generalist insect herbivores and plant defense. Trends Plant Sci 17:293–302 doi:10.1016/j.tplants.2012.02.006

Ahmad M, Sayyed AH, Saleem MA, Ahmad M (2008) Evidence for field evolved resistance to newer insecticides in *Spodoptera litura* (Lepidoptera: Noctuidae) from Pakistan. Crop Prot 27:1367–1372 doi:10.1016/j.cropro.2008.05.003

Ali M, Luttrell R, Young III S (2006) Susceptibilities of *Helicoverpa zea* and *Heliothis virescens* (Lepidoptera: Noctuidae) populations to Cry1Ac insecticidal protein. J Econ Entomol 99:164–175

Allmann S et al. (2013) Feeding-induced rearrangement of green leaf volatiles reduces moth oviposition eLife 2:e00421

Bakkali F, Averbeck S, Averbeck D, Idaomar M (2008) Biological effects of essential oils–a review Food and chemical toxicology 46:446–475

Baldwin IT (2010) Plant volatiles. Current Biology 20:R392–R397 doi:10.1016/j.cub.2010.02.052

Beale MH et al. (2006) Aphid alarm pheromone produced by transgenic plants affects aphid and parasitoid behavior. Proceedings of the National Academy of Sciences 103:10509–10513 doi:10.1073/pnas.0603998103

Bernasconi ML, Turlings TCJ, Ambrosetti L, Bassetti P, Dorn S (1998) Herbivore-induced emissions of maize volatiles repel the corn leaf aphid, *Rhopalosiphum maidis*. Entomologia Experimentalis Et Applicata 87:133–142 doi:DOI 10.1046/j.1570-7458.1998.00315.x

Birkett M, Chamberlain K, Guerrieri E, Pickett J, Wadhams L, Yasuda T (2003) Volatiles from whitefly-infested plants elicit a host-locating response in the parasitoid, *Encarsia formosa*. Journal of chemical ecology 29:1589–1600

Brewer MJ, Trumble JT, Alvarado-Rodriguez B, Chaney WE (1990) Beet armyworm (Lepidoptera: Noctuidae) adult and larval susceptibility to three insecticides in managed habitats and relationship to laboratory selection for resistance Journal of Economic Entomology 83:2136–2146

Brown AWA (1958) Insecticide resistance in arthropods Insecticide resistance in arthropods CABI American cotton bollworm (*Helicoverpa zea*). https://www.cabi.org/isc/datasheet/26776. Accessed November 26 2018

CABI Fall armyworm (*Spodoptera frugiperda*). https://www.plantwise.org/KnowledgeBank/Datasheet.aspx?dsid=29810. Accessed November 26 2018

Capinera JL (1999a) Beet Armyworm; *Spodoptera exigua* (Hübner) (Insecta: Lepidoptera: Noctuidae) University of Florida. http://entnemdept.ufl.edu/creatures/veg/leaf/beet_armyworm.htm. Accessed November 26 2018

Capinera JL (1999b) Cabbage lopper; *Trichoplusia ni* (Hübner) (Insecta: Lepidoptera: Noctuidae). http://entnemdept.ufl.edu/creatures/veg/leaf/cabbage_looper.htm. Accessed November 26 2018

Capinera JL (1999c) Fall armyworm; *Spodoptera frugiperda* (Insecta: Lepidoptera: Noctuidae). http://entnemdept.ufl.edu/creatures/field/fall_armyworm.htm. Accessed November 26 2018

Capinera JL (2001) Tobacco budworm; *Heliothis virescens* (Fabricius) (Insecta: Lepidoptera: Noctuidae). http://entnemdept.ufl.edu/creatures/field/tobacco_budworm.htm. Accessed November 26 2018

Che W, Shi T, Wu Y, Yang Y (2013) Insecticide resistance status of field populations of *Spodoptera exigua* (Lepidoptera: Noctuidae) from China. Journal of Economic Entomology 106:1855–1862 doi:10.1603/EC13128

Christensen SA et al. (2013) The maize lipoxygenase, ZmLOX10, mediates green leaf volatile, jasmonate and herbivore-induced plant volatile production for defense against insect attack The Plant Journal 74:59–73 doi:doi:10.1111/tpj.12101

Cimino AM, Boyles AL, Thayer KA, Perry MJ (2016) Effects of neonicotinoid pesticide exposure on human health: a systematic review Environmental health perspectives 125:155–162

Cna’ani A, Seifan M, Tzin V (2018) Indole is an essential molecule for plant interactions with herbivores and pollinators. J Plant Biol Crop Res 1:1–5

Cordero RJ et al. (2006) Field efficacy of insecticides for control of lepidopteran pests on collards in Virginia Plant health progress 7:32

Degen T, Bakalovic N, Bergvinson D, Turlings TCJ (2012) Differential Performance and Parasitism of Caterpillars on Maize Inbred Lines with Distinctly Different Herbivore-Induced Volatile Emissions PLOS ONE 7:e47589 doi:10.1371/journal.pone.0047589

Dicke M (1986) Volatile spider-mite pheromone and host-plant kairomone, involved in spaced-out gregariousness in the spider mite *Tetranychus urticae*. Physiological Entomology 11:251–262

Dicke M, Sabelis MW (1988) How plants obtain predatory mites as bodyguards. Neth J Zool 38:148–165

Ecobichon DJ (2001) Pesticide use in developing countries Toxicology 160:27–33 doi:https://doi.org/10.1016/S0300-483X(00)00452-2

El-Zaeddi H, Martínez-Tomé J, Calín-Sánchez Á, Burló F, Carbonell-Barrachina ÁA (2016) Volatile Composition of Essential Oils from Different Aromatic Herbs Grown in Mediterranean Regions of Spain Foods (Basel, Switzerland) 5:41 doi:10.3390/foods5020041

Elzen G (1997) Changes in resistance to insecticides in tobacco budworm populations in Mississippi, 1993-1995. The Southwestern entomologist (USA) 22:61–72

Endara M-J et al. (2017) Coevolutionary arms race versus host defense chase in a tropical herbivore–plant system Proceedings of the National Academy of Sciences 114:E7499–E7505 doi:10.1073/pnas.1707727114

Engelberth J, Alborn HT, Schmelz EA, Tumlinson JH (2004) Airborne signals prime plants against insect herbivore attack Proceedings of the National Academy of Sciences 101:1781–1785 doi:10.1073/pnas.0308037100

Erb M, Veyrat N, Robert CA, Xu H, Frey M, Ton J, Turlings TC (2015) Indole is an essential herbivore-induced volatile priming signal in maize Nature communications 6:6273

Fatouros NE, Dicke M, Mumm R, Meiners T, Hilker M (2008) Foraging behavior of egg parasitoids exploiting chemical information. Behav Ecol 19:677–689 doi:10.1093/beheco/arn011

Feeny P (1976) Plant Apparency and Chemical Defense. In: Wallace JW, Mansell RL (eds) Biochemical Interaction Between Plants and Insects. Springer US, Boston, MA, pp 1–40. doi:10.1007/978-1-4684-2646-5_1

Forister ML et al. (2015) The global distribution of diet breadth in insect herbivores Proceedings of the National Academy of Sciences 112:442–447 doi:10.1073/pnas.1423042112

Frost CJ, Appel HM, Carlson JE, De Moraes CM, Mescher MC, Schultz JC (2007) Within-plant signalling via volatiles overcomes vascular constraints on systemic signalling and primes responses against herbivores. Ecol Lett 10:490–498 doi:10.1111/j.1461-0248.2007.01043.x

Frost CJ, Mescher MC, Carlson JE, De Moraes CM (2008a) Plant defense priming against herbivores: getting ready for a different battle. Plant physiol 146:818–824

Frost CJ, Mescher MC, Dervinis C, Davis JM, Carlson JE, De Moraes CM (2008b) Priming defense genes and metabolites in hybrid poplar by the green leaf volatile cis-3-hexenyl acetate. New Phytol 180:722–734 doi:10.1111/j.1469-8137.2008.02599.x

Gasmi L et al. (2018) Can herbivore-induced volatiles protect plants by increasing the herbivores’ susceptibility to natural pathogens? bioRxiv doi:10.1101/317560

Georghiou G (1990) Overview of Insecticide Resistance. In: Managing Resistance to Agrochemicals, vol 421. ACS, pp 18–41

Greenberg SM, Sappington TW, Legaspi BC, Liu TX, Setamou M (2001) Feeding and life history of *Spodoptera exigua* (Lepidoptera: Noctuidae) on different host plants. Ann Entomol Soc Am 94:566–575 doi:Doi 10.1603/0013-8746(2001)094[0566:Falhos]2.0.Co;2

Hahn M, Schotthöfer A, Schmitz J, Franke LA, Brühl CA (2015) The effects of agrochemicals on Lepidoptera, with a focus on moths, and their pollination service in field margin habitats Agriculture, Ecosystems & Environment 207:153–162

Hardee D, Adams L, Elzen G (2001) Monitoring for changes in tolerance and resistance to insecticides in bollworm/tobacco budworm in Mississippi, 1996-1999. Southwestern Entomologist 26:365–372

Harding JA (1976) Heliothis spp.: seasonal occurrence, hosts and host importance in the lower Rio Grande Valley. Environ Entomol 5:666–668

Hardke JT, Temple JH, Leonard BR, Jackson RE (2011) Laboratory toxicity and field efficacy of selected insecticides against fall armyworm (Lepidoptera: Noctuidae). Fla Entomol 94:272–278

Hare JD (2011) Ecological role of volatiles produced by plants in response to damage by herbivorous insects Annual Review of Entomology 56:161–180

Heil M (2004) Direct defense or ecological costs: responses of herbivorous beetles to volatiles released by wild lima bean (*Phaseolus lunatus*). Journal of chemical ecology 30:1289–1295

Heil M (2014) Herbivore-induced plant volatiles: targets, perception and unanswered questions New Phytologist 204:297–306 doi:10.1111/nph.12977

Heil M, Bueno JCS (2007) Within-plant signaling by volatiles leads to induction and priming of an indirect plant defense in nature. Proceedings of the National Academy of Sciences 104:5467–5472

Hoo CS, Coudriet D, Vail P (1984) *Trichoplusia ni* (Lepidoptera: Noctuidae) larval development on wild and cultivated plants. Environ Entomol 13:843–846

Howard V. C., Bradford A. H. (2003) Herbivore Responses to Plant Secondary Compounds: A Test of Phytochemical Coevolution Theory The American Naturalist 161:507–522 doi:10.1086/368346

Hubert J, Münzbergová Z, Santino A (2008) Plant volatile aldehydes as natural insecticides against stored-product beetles. Pest Management Science 64:57–64 doi:doi:10.1002/ps.1471

Huffaker A et al. (2013) Plant elicitor peptides are conserved signals regulating direct and indirect antiherbivore defense Proceedings of the National Academy of Sciences 110:5707–5712 doi:10.1073/pnas.1214668110

Isman MB (2016) Pesticides based on plant essential oils: phytochemical and practical considerations. In: Medicinal and aromatic crops: production, phytochemistry, and utilization, vol 1218. ACS Symposium Series, vol 1218. American Chemical Society, pp 13–26. doi:10.1021/bk-2016-1218.ch002

Jurriaan T et al. (2007) Priming by airborne signals boosts direct and indirect resistance in maize The Plant Journal 49:16–26 doi:doi:10.1111/j.1365-313X.2006.02935.x

Kessler A, Baldwin IT (2001) Defensive function of herbivore-induced plant volatile emissions in nature. Science 291:2141–2144 doi:10.1126/science.291.5511.2141

Krieger RI, Feeny PP, Wilkinson CF (1971) Detoxication enzymes in the guts of caterpillars: an evolutionary answer to plant defenses? Science 172:579–581

Laquale S, Avato P, Argentieri MP, Bellardi MG, D’Addabbo T (2018) Nematotoxic activity of essential oils from Monarda species. Journal of Pest Science doi:10.1007/s10340-018-0957-1

Lee J-H, Wood TK, Lee J (2015) Roles of indole as an interspecies and interkingdom signaling molecule. Trends in Microbiology 23:707–718 doi:10.1016/j.tim.2015.08.001

Lee S, Tsao R, Coats JR (1999) Influence of dietary applied monoterpenoids and derivatives on survival and growth of the european corn borer (lepidoptera: pyralidae). Journal of Economic Entomology 92:56–67 doi:10.1093/jee/92.1.56

Liao C, Heckel DG, Akhurst R (2002) Toxicity of *Bacillus thuringiensis* insecticidal proteins for *Helicoverpa armigera* and *Helicoverpa punctigera* (Lepidoptera: Noctuidae), major pests of cotton. Journal of Invertebrate Pathology 80:55–63 doi:10.1016/S0022-2011(02)00035-6

Liburd O, Funderburk J, Olson S (2000) Effect of biological and chemical insecticides on Spodoptera species (Lep., Noctuidae) and marketable yields of tomatoes Journal of applied entomology 124:19–25

Loreto F, Ciccioli P, Brancaleoni E, Cecinato A, Frattoni M (1998) Measurement of isoprenoid content in leaves of Mediterranean Quercus spp. by a novel and sensitive method and estimation of the isoprenoid partition between liquid and gas phase inside the leaves Plant Science 136:25–30 doi:https://doi.org/10.1016/S0168-9452(98)00092-2

Loreto F, Nascetti P, Graverini A, Mannozzi M (2000) Emission and content of monoterpenes in intact and wounded needles of the Mediterranean Pine, *Pinus pinea*. Funct Eol 14:589–595 doi:doi:10.1046/j.1365-2435.2000.t01-1-00457.x

Maffei ME, Gertsch J, Appendino G (2011) Plant volatiles: production, function and pharmacology Natural product reports 28:1359–1380

Martin P, Lingren P, Greene G (1976) Relative abundance and host preferences of cabbage looper, soybean looper, tobacco budworm, and corn earworm on crops grown in northern Florida. Environ Entomol 5:878–882

McEwen F, Splittstoesser C (1970) Resistance to organophosphate insecticides in the cabbage looper in New York. Journal of Economic Entomology 63

Moar WJ, Trumble JT, Federici BA (1989) Comparative toxicity of spores and crystals from the nrd-12 and hd-1 strains of *Bacillus thuringiensis* suhsp. kurstaki to neonate beet armyworm (lepidoptera: noctuidae). Journal of Economic Entomology 82:1593–1603 doi:10.1093/jee/82.6.1593

Moghaddam M, Mehdizadeh L (2017) Chemistry of Essential Oils and Factors Influencing Their Constituents. In: Soft Chemistry and Food Fermentation. Elsevier, pp 379–419

Monson RK, Grote R, Niinemets Ü, Schnitzler JP (2012) Modeling the isoprene emission rate from leaves. New Phytol 195:541–559 doi:10.1111/j.1469-8137.2012.04204.x

Mossa A-TH(2016) Green pesticides: Essential oils as biopesticides in insect-pest management. Journal of Environmental Science and Technology 9:354

Mulé R, Sabella G, Robba L, Manachini B (2017) Systematic Review of the Effects of Chemical Insecticides on Four Common Butterfly Families Frontiers in Environmental Science 5 doi:10.3389/fenvs.2017.00032

Niinemets Ü, Kännaste A, Copolovici L (2013) Quantitative patterns between plant volatile emissions induced by biotic stresses and the degree of damage. Front plant sci 4 doi:10.3389/fpls.2013.00262

Niinemets Ü, Loreto F, Reichstein M (2004) Physiological and physicochemical controls on foliar volatile organic compound emissions Trends in Plant Science 9:180–186 doi: https://doi.org/10.1016/j.tplants.2004.02.006

Niu Y, Meagher Jr RL, Yang F, Huang F (2013) Susceptibility of field populations of the fall armyworm (Lepidoptera: Noctuidae) from Florida and Puerto Rico to purified Cry1F protein and corn leaf tissue containing single and pyramided Bt genes. Fla Entomol 96:701–713

Ormeño E, Goldstein A, Niinemets Ü (2011) Extracting and trapping biogenic volatile organic compounds stored in plant species. Trends in Analytical Chemistry 30:978–989 doi:10.1016/j.trac.2011.04.006

Pearson AC (1983) Biology, population dynamics, and pest status of the beet armyworm (Spodoptera exigua) in the Imperial Valley of California

Pichersky E, Gershenzon J (2002) The formation and function of plant volatiles: perfumes for pollinator attraction and defense. Current Opinion in Plant Biology 5:237–243 doi:10.1016/S1369-5266(02)00251-0

Pimentel D (1996) Green revolution agriculture and chemical hazards Science of The Total Environment 188:S86–S98 doi:https://doi.org/10.1016/0048-9697(96)05280-1

R (2018) A language and environment for statistical computing R foundation for statistical computing, Vienna, Austria

Rivera-Vega LJ, Galbraith DA, Grozinger CM, Felton GW (2017) Host plant driven transcriptome plasticity in the salivary glands of the cabbage looper (*Trichoplusia ni*). PLoS ONE 12:e0182636 doi:10.1371/journal.pone.0182636

Sadeghi A, Van Damme EJM, Smagghe G (2009) Evaluation of the susceptibility of the pea aphid, *Acyrthosiphon pisum*, to a selection of novel biorational insecticides using an artificial diet. J Insect Sci 9:1–8 doi:10.1673/031.009.6501

Sandra I et al. (2014) Herbivore-induced poplar cytochrome P450 enzymes of the CYP71 family convert aldoximes to nitriles which repel a generalist caterpillar. Plant J 80:1095–1107 doi:doi:10.1111/tpj.12711

Schiestl FP, Ayasse M (2001) Post-pollination emission of a repellent compound in a sexually deceptive orchid: a new mechanism for maximising reproductive success? Oecologia 126:531–534 doi:10.1007/s004420000552

Schiestl FP, Ayasse M, Paulus HF, Löfstedt C, Hansson BS, Ibarra F, Francke W (1999) Orchid pollination by sexual swindle. Nature 399:421 doi:10.1038/20829

Schmelz EA, Alborn HT, Banchio E, Tumlinson JH (2003) Quantitative relationships between induced jasmonic acid levels and volatile emission in Zea mays during Spodoptera exigua herbivory Planta 216:665–673 doi:10.1007/s00425-002-0898-y

Schnee C, Köllner TG, Held M, Turlings TC, Gershenzon J, Degenhardt J (2006) The products of a single maize sesquiterpene synthase form a volatile defense signal that attracts natural enemies of maize herbivores. Proc Nat Acad Sci USA 103:1129–1134

Shiojiri K et al. (2006) Changing green leaf volatile biosynthesis in plants: An approach for improving plant resistance against both herbivores and pathogens. Proc Nat Acad Sci USA 103:16672–16676 doi:10.1073/pnas.0607780103

Slansky F (1993) Xanthine toxicity to caterpillars synergized by allopurinol, a xanthine dehydrogenase/oxidase inhibitor. Journal of chemical ecology 19:2635–2650

Sparks TC, Nauen R (2015) IRAC: Mode of action classification and insecticide resistance management Pesticide Biochemistry and Physiology 121:122–128 doi:https://doi.org/10.1016/j.pestbp.2014.11.014

Sugimoto K, Matsui K, Takabayashi J (2015) Conversion of volatile alcohols into their glucosides in Arabidopsis. Communicative & Integrative Biology 8:e992731 doi:10.4161/19420889.2014.992731

Tingle CC, Rother JA, Dewhurst CF, Lauer S, King WJ (2003) Fipronil: environmental fate, ecotoxicology, and human health concerns. In: Reviews of environmental contamination and toxicology. Springer, pp 1–66

Tominaga T, Dubourdieu D (2000) Identification of cysteinylated aroma precursors of certain volatile thiols in passion fruit juice. Journal of Agricultural and Food Chemistry 48:2874–2876 doi:10.1021/jf990980a

Turlings TC, Loughrin JH, McCall PJ, Röse US, Lewis WJ, Tumlinson JH (1995) How caterpillar-damaged plants protect themselves by attracting parasitic wasps. Proc Nat Acad Sci USA 92:4169–4174

Turlings TC, Tumlinson JH, Lewis WJ (1990) Exploitation of herbivore-induced plant odors by host-seeking parasitic wasps. Science 250:1251–1253

Unsicker SB, Kunert G, Gershenzon J (2009) Protective perfumes: the role of vegetative volatiles in plant defense against herbivores. Current Opinion in Plant Biology 12:479–485 doi:10.1016/j.pbi.2009.04.001

Valenta K, Nevo O, Martel C, Chapman CA (2017) Plant attractants: integrating insights from pollination and seed dispersal ecology. Evol Biol 31:249–267 doi:10.1007/s10682-016-9870-3

Veyrat N, Robert CAM, Turlings TCJ, Erb M (2016) Herbivore intoxication as a potential primary function of an inducible volatile plant signal. Journal of Ecology 104:591–600

Vickers CE, Gershenzon J, Lerdau MT, Loreto F (2009) A unified mechanism of action for volatile isoprenoids in plant abiotic stress. Nat Chem Biol 5:283 doi:10.1038/nchembio.158

von Mérey G, Veyrat N, D’Alessandro M, Turlings T (2013) Herbivore-induced maize leaf volatiles affect attraction and feeding behavior of Spodoptera littoralis caterpillars. Front Plant Sci 4 doi:10.3389/fpls.2013.00209

Vreysen MJ, Klassen W, Carpenter JE (2016) Overview of technological advances toward greater efficiency and efficacy in sterile insect-inherited sterility programs against moth pests Florida Entomologist 99:1–13

Waters DJ, Barfield CS (1989) Larval development and consumption by *Anticarsia gemmatalis* (Lepidoptera: Noctuidae) fed various legume species. Environ Entomol 18:1006–1010

Whittaker RH, Feeny PP (1971) Allelochemics: chemical interactions between species. Science 171:757–770

Wickham H (2011) ggplot2: Elegant Graphics for Data Analysis. J R Stat Soc a Stat doi DOI:10.1111/j.1467-985X.2010.00676_9.x

Ye M, Glauser G, Lou Y, Erb M, Hu L (2019) Molecular dissection of early defense signaling underlying volatile-mediated defense regulation and herbivore resistance in rice The Plant Cell:tpc.00569.02018 doi:10.1105/tpc.18.00569

Yi H-S, Heil M, Adame-Álvarez RM, Ballhorn DJ, Ryu C-M (2009) Airborne induction and priming of plant defenses against a bacterial pathogen. Plant Physiol 151:2152–2161 doi:10.1104/pp.109.144782

Yu SJ, Nguyen SN, Abo-Elghar GE (2003) Biochemical characteristics of insecticide resistance in the fall armyworm, *Spodoptera frugiperda* (J.E. Smith). Pestic Biochem Physiol 77:1–11 doi:10.1016/S0048-3575(03)00079-8

Zakir A, Bengtsson M, Sadek MM, Hansson BS, Witzgall P, Anderson P (2013) Specific response to herbivore-induced de novo synthesized plant volatiles provides reliable information for host plant selection in a moth. J Exp Biol 216:3257–3263 doi:10.1242/jeb.083188

Zalucki MP, Clarke AR, Malcolm SB (2002) Ecology and behavior of first instar larval Lepidoptera Annual Review of Entomology 47:361–393

Zhao Y et al. (2017) Effects of the plant volatile trans‑2-hexenal on the dispersal ability, nutrient metabolism and enzymatic activities of *Bursaphelenchus xylophilus*. Pestic Biochem Physiol doi:10.1016/j.pestbp.2017.08.004

Zhu J, Park KC (2005) Methyl salicylate, a soybean aphid-induced plant volatile attractive to the predator *Coccinella septempunctata*. Journal of chemical ecology 31:1733–1746

